# A prognostic matrix code defines functional glioblastoma phenotypes and niches

**DOI:** 10.1101/2023.06.06.543903

**Authors:** Monika Vishnoi, Zeynep Dereli, Zheng Yin, Elisabeth K. Kong, Meric Kinali, Kisan Thapa, Ozgun Babur, Kyuson Yun, Nourhan Abdelfattah, Xubin Li, Behnaz Bozorgui, Robert C. Rostomily, Anil Korkut

## Abstract

Interactions among tumor, immune and vascular niches play major roles in driving glioblastoma (GBM) malignancy and treatment responses. The composition, heterogeneity, and localization of extracellular core matrix proteins (CMPs) that mediate such interactions, however, are not well understood. Here, we characterize functional and clinical relevance of genes encoding CMPs in GBM at bulk, single cell, and spatial anatomical resolution. We identify a “matrix code” for genes encoding CMPs whose expression levels categorize GBM tumors into matrisome-high and matrisome-low groups that correlate with worse and better survival, respectively, of patients. The matrisome enrichment is associated with specific driver oncogenic alterations, mesenchymal state, infiltration of pro-tumor immune cells and immune checkpoint gene expression. Anatomical and single cell transcriptome analyses indicate that matrisome gene expression is enriched in vascular and leading edge/infiltrative anatomic structures that are known to harbor glioma stem cells driving GBM progression. Finally, we identified a 17-gene matrisome signature that retains and further refines the prognostic value of genes encoding CMPs and, importantly, potentially predicts responses to PD1 blockade in clinical trials for GBM. The matrisome gene expression profiles may provide biomarkers of functionally relevant GBM niches that contribute to mesenchymal-immune cross talk and patient stratification to optimize treatment responses.

## Introduction

Glioblastoma (GBM) is an aggressive disease and meaningful improvements in survival have yet to be realized in a significant number of patients {Taslimi, 2021 #314}. A high degree of intratumor heterogeneity, invasive growth and myriad mechanisms of treatment resistance limit the effectiveness of standard of care chemoradiation, targeted precision approaches and immunotherapy {Kumar, 2014 #162; Lee, 2017 #84}. Central to the success of precision therapy is the integration of molecular and/or cellular data that accurately capture therapeutic targets and prognostic markers. Aside from the *MGMT* promoter methylation status for guiding temozolomide-based treatments, actionable driver mutations and immune checkpoint molecule expression have emerged as candidate biomarkers to inform targeted therapy and immunotherapy, respectively, for managing GBM {Tan, 2020 #319}. Despite their conceptual appeal, precision therapy and immunotherapy have yet to significantly improve durable outcomes {Gilbert, 2013 #316; Lim, 2022 #313; Stupp, 2005 #315; Stupp, 2017 #281; Stupp, 2015 #282; Taslimi, 2021 #314; Weller, 2016 #317; Wick, 2017 #318}. The lack of progress with these approaches, and limited efficacy of standard genotoxic therapies, reflects, in part, insufficient understanding of factors that impact GBM malignancy and treatment responses.

The matrisome is the collection of extracellular matrix (ECM) proteins {Naba, 2012 #97; Hynes, 2012 #98}. It is comprised of the core matrix proteins (glycoproteins, collagens, and proteoglycans), and matrisome associated proteins (secreted factors, regulators, and ECM affiliated proteins) {Naba, 2016 #87; Naba, 2012 #97}. The matrisome plays critical roles in regulating normal and pathologic processes, including cancer malignancy {Karamanos, 2021 #439} {Rafaeva, 2020 #433}. Mechanistically, matrisome-tumor interactions contribute to cancer phenotypes through ligand-receptor interactions or biophysical structural effects that directly regulate tumor and stromal cell signaling or indirectly through modulating the tumor microenvironment (TME) {Henke, 2019 #56}. Of relevance to precision oncology, matrisome-cancer cell interactions are likely to modify drug responses predicted solely based on mutation profiles. A deeper understanding of the matrisome-GBM interactome is expected to refine the performance of *ex vivo* organotypic and 3D pre-clinical GBM models to predict treatment responses and inform novel mechanistic interactions that underlie GBM malignancy {Rodriguez, 2020 #84; Horowitz, 2020 #83, Wan, 2020 #480;Sood, 2019 #467;Neufeld, 2022 #477;Gomez-Roman, 2017 #468;Cornelison, 2022 #469;Neufeld, 2021 #470}. While the genomic landscape in GBM has been well characterized {Cancer Genome Atlas Research, 2008 #48} {Puchalski, 2018 #49} {Patel, 2014 #75}, the functional and clinical importance as well as the diversity of the GBM matrisome have not been well established {Langlois, 2014 #331; Laurentino, 2022 #329; Sethi, 2022 #330}.

To achieve these goals, we undertook a comprehensive analysis of the genes encoding core matrix proteins (CMPs) in GBM. Through analysis of the expression of these genes in the TCGA GBM dataset, we identified three groups of GBM with matrix-high (M-H), matrix-low type a (M-La) and matrix-low type b (M-Lb) gene expression profiles. Importantly, the M-H profile predicts worse clinical outcomes and is associated with oncogenic processes, including epithelial mesenchymal transition (EMT), a pro-tumor immune signature and signaling relevant to GBM malignancy. Using the IVYGap GBM database, we detected enrichment of the CMP-encoding gene expression signatures in anatomic sub-regions such as vascular and infiltrative regions, suggesting that the CMP profile may comprise a “code” that defines functional GBM niches. Consistent with this, single cell RNA expression analysis revealed enrichment of CMPs primarily in pericytes and endothelial cells with moderate expression in glioma cells and sparse expression in immune cells. Finally, we identified a 17-gene matrisome signature that can potentially predict both survival outcomes and response to anti-PD1 blockade in GBM. Together these results provide evidence that CMP composition, localization and heterogeneity can mediate malignant phenotypes, clinical outcomes, and treatment responses in GBM.

## Results

### A compendium of multi-modal molecular and clinical data from GBM patients

To comprehensively characterize the GBM matrisome, we established a compendium of genomic, transcriptomic, and phosphoproteomic datasets. In our analysis, we included the core matrisome proteins (CMPs) glycoproteins, collagens, and proteoglycans (n=276) {Naba, 2012 #97; Naba, 2017 #83; Naba, 2016 #87} (**Supplementary Table 1**). To characterize the interpatient matrisome heterogeneity, we included 157 IDH wild type (WT) GBM samples from 151 patients with varying coverages for genomic, transcriptomic, proteomic, and clinical data from the TCGA GBM dataset {Cancer Genome Atlas Research, 2008 #122} (**Supplementary Table 2)**. We mapped the spatio-anatomical heterogeneity of matrisome using the transcriptomic data from the IvyGap repository (245 samples across 7 anatomic regions in 34 tumors) {Puchalski, 2018 #260}. The matrisome heterogeneity across tumor niches was further characterized using single cell transcriptomics data from 201,986 glioma, immune, and other stromal cells in 16 IDH WT GBM and 2 IDH mutant low-grade glioma tumors from patients {Abdelfattah, 2022 #100}. The proteomic outcomes of the transcriptomic signatures were determined using mass-spectroscopy data and RNA sequencing data available for IDH-wild type GBM tumors (N=86) in the CPTAC repository. For each data modality, we also integrated the relevant clinical and phenotypic parameters to establish multi-faceted interactions between the ECM and key GBM phenotypes, including immune, vascularization, cell signaling, and differentiation (e.g., EMT) states. We studied the therapeutic implications of the matrisome enrichment in GBM using transcriptomic and survival data from a clinical trial of 28 patients with resectable recurrent GBM treated with anti-PD1 therapy {Cloughesy, 2019 #351}. The patients (N=164) with resectable and recurrent GBM, who did not receive anti-PD1 therapy served as the control cohort to distinguish the impact of anti-PD1 on patients with different matrisome states. The resulting multi-modal data compendium enabled us to establish a robust matrisome code that predicts patient survival, disease characteristics and therapy responses in GBM (**Fig. 1A**).

**Figure 1.**
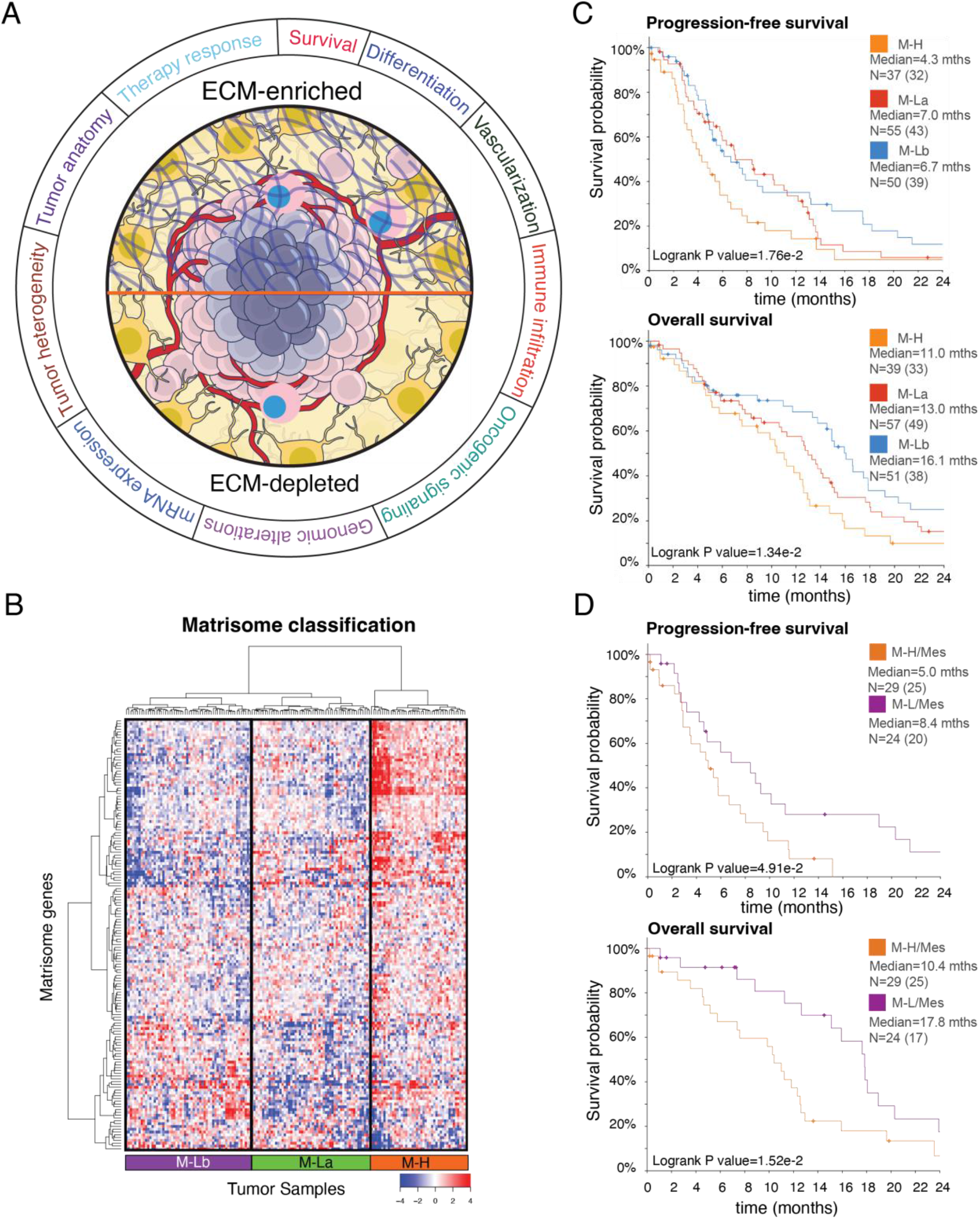
The landscape of ECM composition in GBM and patient survival. ***A.** The association of extracellular matrix composition with key molecular and physiological states in GBM. **B.** The hierarchical clustering of mRNA levels of core extracellular matrix genes (matrisome, N=274) in 157 GBM tumors from 151 patients (source: TCGA, Level 3 mRNA sequencing data). The genes with low expression variation (σ < 1) across the tumors are filtered out. The final set included 154 genes that are most variant across samples. Three patient groups are identified as Matrisome high (M-H), Matrisome Low A (M-La) and Matrisome Low B (M-Lb) based on overall matrisome expression. **C.** The progression-free survival and overall survival of patients with M-H, M-La, and M-Lb GBM. N is the number of patients. **D.** The progression-free and overall survival of patients with mesenchymal transcriptional subtype and M-H or M-L matrisome subtype of GBM. The M-L cohort includes both M-La and M-Lb. In all survival curves, the number of events (death or disease progression) is given in parentheses. The marks on survival curves represent censored data points*. mths=months. The survival curves are truncated at 24 months (**Supplementary Fig. 1B for complete curves)**.

### Core matrix gene expression signatures predict GBM patient survival

To investigate whether the interpatient heterogeneity of the core matrisome defines clinically relevant subgroups of GBM, we analyzed the distribution of CMP-encoding gene expression levels in the TCGA GBM dataset {Brennan, 2009 #116}. Unsupervised clustering analysis identified three major clusters of GBM with different CMP-encoding gene expression levels (**Fig. 1B**), which were termed matrix-high (M-H, n=44 cases, 28% of the cohort), matrix-low type a (M-La, n=58 cases, 37% of the cohort) and matrix-low type b (M-Lb, n=55 cases, 35% of the cohort) (**Fig. 1B, Table 1).** Compared to the M-H sub-group, overall CMP-encoding gene expression levels were lower in the two M-L subgroups, M-La and M-Lb. These CMP sub-groups correlated with differential progression-free survival (PFS) and overall survival (OS), with M-H having the worst OS and PFS (Median PFS (months): M-H= 4.3, M-La= 7.0, M-Lb=6.7, log-rank p = 1.76E-2; Median OS (months): M-H= 11.0, M-La= 13.0, M-Lb=16.1, log-rank p = 1.34E-2) (**Fig. 1C and Supplementary Table 1**). When M-La and M-Lb cohorts are combined into a single matrix-low group (M-L) and compared to M-H, the significant differences in PFS and OS remained (log rank p= 6.01E-3 and 1.13E-2, respectively) (**Supplementary Fig. 1**). In a multivariate analysis accounting for gender, age, GBM transcriptomic subtypes, MGMT methylation status and treatment (chemoradiation vs radiation alone); M-H versus M-La or M-Lb independently predicted shorter survivals (p= 3.0E-3 and 2.7E-13, respectively; **Supplementary Table 3**).

**Table 1.** The demographic profile of TCGA sample cohort.

|  |  | M-H (44) | M-La (58) | M-Lb (55) | Total | P-value |
| --- | --- | --- | --- | --- | --- | --- |
| **Gender** | Male | 28 | 42 | 31 | 101 | 0.084* |
|  | Female | 16 | 15 | 24 | 55 | 0.117* |
|  | Not available | 0 | 1 | 0 | 1 | NA |
| **Age Median(Std dev)** | Male | 59±10.62 | 59±13.71 | 63±9.78 |  | 0.294** |
|  | Female | 63±12.26 | 62±15.62 | 60±10.34 |  | 0.743** |
| **Karnofsky Performance Score** | M-H=29; M-La=42; M-Lb=46 | 80±12.45 | 80±14.13 | 80±15.27 |  | 1.000** |
| **Vital status** | Living | 11 | 19 | 20 | 50 | 0.473* |
|  | Deceased | 33 | 38 | 35 | 106 | 0.259* |
|  | Not available | 0 | 1 | 0 | 1 | NA |
| **Gene Expression Subtype** | Proneural | 3 | 21 | 5 | 29 | < 1e-04* |
|  | Neural | 4 | 14 | 9 | 27 | 0.085* |
|  | Classical | 5 | 13 | 23 | 41 | 0.002* |
|  | Mesenchymal | 30 | 8 | 17 | 55 | < 1e-04* |
|  | G-CIMP | 0 | 1 | 0 | 1 | NA |
|  | Not available | 2 | 1 | 1 | 4 | NA |
| **Treatment** | Chemo-radiation | 26 | 39 | 34 | 99 | 0.494* |
|  | Chemotherapy only | 0 | 0 | 1 | 1 | NA |
|  | Radiation only | 15 | 18 | 19 | 52 | 0.859* |
|  | Other (Unspecified + Not Available) | 3 | 1 | 1 | 5 | NA |
| **MGMT Status** | Unmethylated | 20 | 25 | 28 | 73 | 0.503* |
|  | Methylated | 11 | 17 | 20 | 48 | 0.278* |
|  | Not available | 13 | 16 | 7 | 36 | 0.290* |

\* Fisher's exact test (see methods). \*\* 1-way ANOVA test.

Consistent with their known prognostic importance, unmethylated MGMT, radiation only versus chemoradiation treatment, and increased age also demonstrated significant negative associations with survival (p=3.33e-15, 6.77e-12, and 1.49e-09, respectively).

We determined that each CMP subgroup is enriched in a specific transcriptional subtype {Verhaak, 2010 #293} (**Table 1**). The M-H subgroup is enriched in the Mesenchymal GBM (mesGBM) subtype (p < 1e-04). On the other hand, the M-Lb subgroup is associated with the Classical GBM subtype (p=2e-03), while the M-La subgroup is enriched in the Proneural subtype (p < 1e-04) and, with a non-significant trend, in the Neural subtype (p=0.085), whose relevance to GBM biology has recently come under suspicion due to potential contamination of the samples from normal neural tissue. Importantly the M-H subgroup also predicted significantly shorter PFS and OS (log-rank p= 4.91e-2 and 1.52e-2, respectively) within the mesGBM subtype (**Fig. 1D**). Despite the differential enrichment of CMP subgroups in each of transcriptional subtypes, the significantly poor survival outcomes of patients with M-H tumors compared to those with M-L tumors within the mesenchymal subtype suggests that the matrisome intertumor heterogeneity and subgroups are clinically relevant and the impact on survival is not driven by the transcriptional subtypes. Together these data indicate that differential expression of CMP genes provide a robust and independent prognostic biomarker of clinical outcomes in IDH WT GBM.

### Matrisome signature expression is associated with expression of mesenchymal and immune markers

We next analyzed molecular correlates of each CMP sub-group. Consistent with the associations between the transcriptomic and matrisome subtypes, the mesenchymal subtype-associated NF1 mutations and classical subtype-associated EGFR alterations, primarily amplifications, are enriched in the M-H and M-Lb subgroups, respectively {Verhaak, 2010 #293} **(Fig. 2A).** No mutations associated with the proneural subtype (e.g., PDGFRα amplification) are enriched in the M-La subgroup (**Fig. 2A**). Gene set enrichment analysis (GSEA) of differentially expressed genes (DEGs) in M-H versus M-L identified the enrichment of transcripts associated with leukocyte migration, leukocyte activation, cell motility, angiogenesis, ECM organization, and cell adhesion in the M-H subgroup (**Fig. 2B, Supplementary Fig. 2**). The phosphoproteomics data (available in 23 M-H and 54 M-L samples for 191 total protein or phosphoprotein markers) enabled differential proteomics and pathway activation analyses (**Fig. 2C**) {Chen, 2019 #483} {Li, 2013 #482}. PAI1 (Serpine 1), Fibronectin (FN1) and Caveolin1 are the most upregulated proteins in M-H while Beta-catenin is significantly enriched in M-L CMP subgroups. PAI1 promotes GBM invasion, is upregulated in mesenchymal GBM subtypes and is associated with shorter overall survival {Seker, 2019 #332}. Increased FN1 and Caveolin1 are also associated with increased GBM malignancy {Kabir, 2022 #353; Wu, 2022 #352} {Pu, 2019 #357; Moriconi, 2021 #355}. Notably the receptor tyrosine kinase (RTK) and core reactive {Ha, 2018 #345} pathways are enriched in M-Lb and M-H, respectively (**Fig. 2C, Supplementary Fig. 2**). The RTK pathway activity in the M-Lb samples is consistent with EGFR amplification enrichment in this sub-group. Activation of the core reactive pathway, comprised of stromal proteins, Claudin7, E-cadherin, Beta-catenin, RBM15 and Caveolin1, was first discovered as associated with poor survival in breast cancers {Ha, 2018 #345}. Finally, consistent with EMT hallmarks of cell motility, angiogenesis, ECM organization and cell adhesion; tumors within the M-H subgroup are enriched with the EMT markers based on both transcriptomics- and proteomics-based scoring of EMT-signature genes {Byers, 2013 #481} {Mak, 2016 #333} {Akbani, 2014 #339} (P values < 1E-4). (**Fig. 2D, Supplementary Fig. 2**). In conclusion, CMP subgroups are enriched in genomic alterations consistent with their associated transcriptional subtypes, and importantly, demonstrate robust correlations with signatures of mesenchymal state, immune activation, and tumor malignancy (e.g., motility, adhesion, angiogenesis).

**Figure 2.**
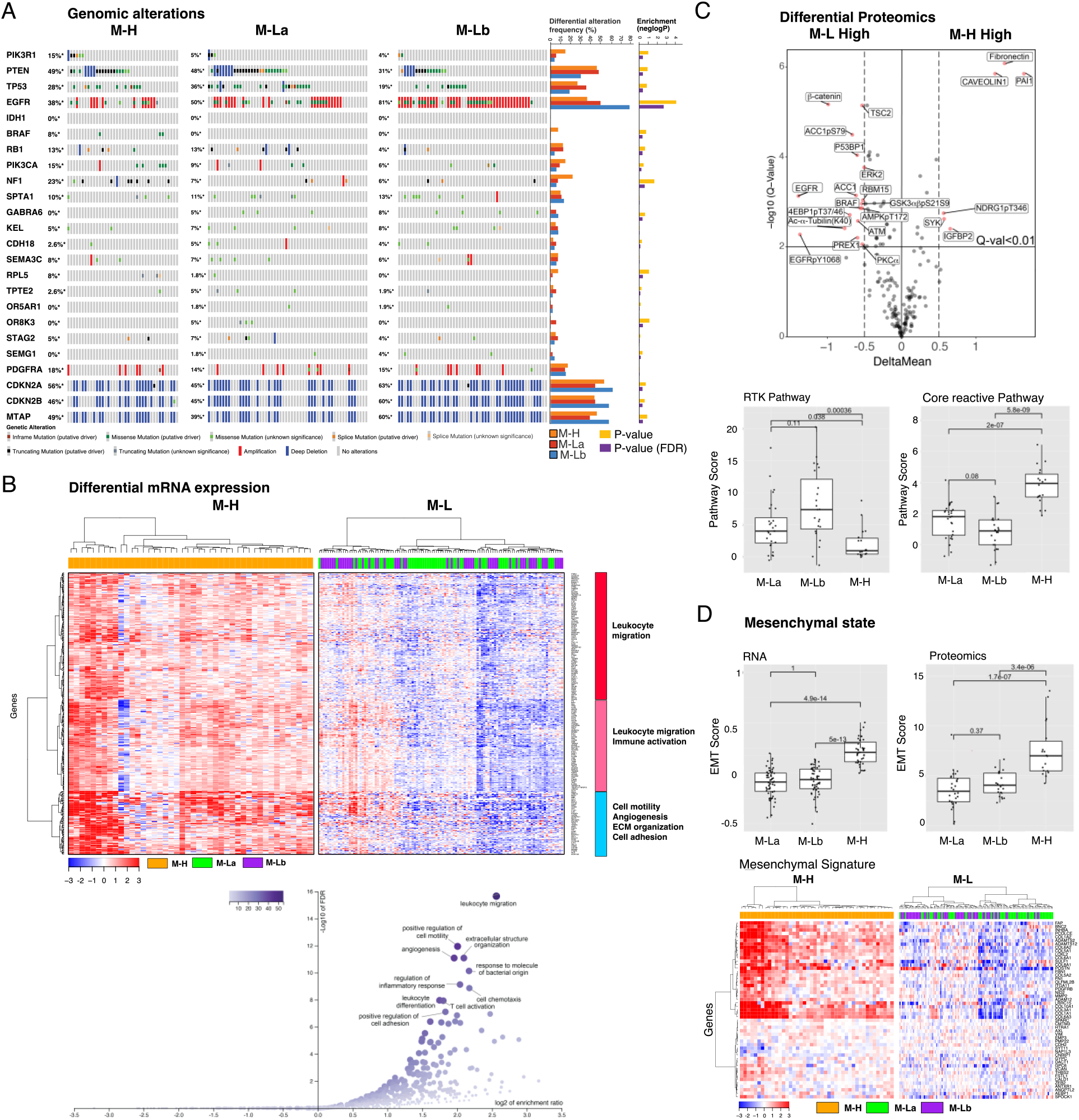
The multi-omic landscape of matrisome-enrichment in GBM. ***A.*** *Mutational and copy-number alterations in GBM stratified by the matrisome subgroups. The p-values are based on Fisher’s exact-test (FDR-adjusted using the Benjamini-Hochberg (BH) method) and quantify the significance of the alteration enrichment across the matrisome groups. **B.** The differential mRNA expression between M-H vs. M-L groups. Genes whose mRNA levels are significantly higher (FDR-adjusted P-values < 0.01 in a non-parametric Wilcoxon-test) in the M-H group are included in the analysis. Three differentially expressed gene groups are defined based on a hierarchical clustering of mRNA expression in the M-H cohort. The functional enrichment within each group is annotated based on a gene set enrichment analysis (Web-Gestalt method). **C.** The differential phosphoprotein and total protein levels in M-H vs. M-L tumors based on reverse-phase protein array (RPPA) profiling of the TCGA cohort (n=77) {Li, 2013 #482} {Chen, 2019 #483}. On the volcano plot, the x-axis is the difference of mean expressions across the samples (<X>_M-H_- <X>_M-L_) and the y-axis is the p-values based on a t-test (adjusted with Bonferroni correction) that quantify the significance of protein expression differences between M-H and M-L (M-La + M-Lb) groups. The boxplots quantify the differential pathway activities. The pathway activity scores are defined as cumulative expression levels of proteins that function in the corresponding pathways as defined in {Akbani, 2014 #339}. The activities of receptor tyrosine kinase (RTK) and core reactive (representing stroma gene expression) pathways are significantly different between the M-H vs M-L groups. **D.** The representation of epithelial vs. mesenchymal differentiation states in M-H vs. M-L groups. The proteomic ({Akbani, 2014 #338} {Akbani, 2014 #339}) and transcriptomic ({Byers, 2013 #481}) markers of EMT are used to calculate the EMT scores as defined in the respective references. The boxplots demonstrate the statistically significant difference between the M-H vs M-L groups (p-values are based on a Wilcoxon-test). The heatmap visualizes the differential mRNA expression of mesenchymal genes in the matrisome groups (see **Supplementary Fig. 2** for the epithelial gene expression signature)*.

### Expression of CMP-encoding genes is associated with a pro-tumor immune infiltration

Based on the observation that immune-related processes were enriched in the M-H subgroup, we analyzed the characteristics of the tumor-immune microenvironment (TIME) across CMP subgroups. We first quantified the overall immune infiltration using a methylation-based immune infiltration score for each tumor sample {Hoadley, 2018 #484} and compared it across CMP subgroups. We observed a significantly higher immune infiltration in the M-H subgroup compared to the M-L subgroup **(Fig. 3A)**. We further assessed the enrichment of specific immune cell types within the TIME by analyzing the RNA-based CIBERSORT deconvolution results available for 157 GBM samples from the TCGA repository (**Fig. 3A**) {Thorsson, 2018, #287}. This analysis identified an enrichment of pro-tumor immune cell types, including M2 macrophages, neutrophils, resting NK cells and regulatory T cells (Tregs) in M-H tumors. Analysis of 31 immune checkpoint (ICP) genes that we curated based on the FDA-approval status and ongoing clinical trials of immunotherapies {Bozorgui, 2021 #505} reveals that expression levels of genes encoding therapeutically actionable immune checkpoints also correlate with CMP subgroups and immune infiltration (**Fig. 3B, Supplementary Fig. 3**). Out of these 31 ICP genes, mRNA levels of 17 genes are significantly higher in the M-H subgroup (**Supplementary Fig. 3**). The difference in the expression of CD276 (B7-H3), an orphan receptor with immune-suppressive, angiogenic and EMT functions, between M-H vs M-L subgroups is most significant. By comparing CMP subgroup-specific expression patterns of known immune receptor-ligand pairs, we mapped putative functional interactions that could be relevant within the GBM tumor-immune interface {Verhaak, 2010 #293}. In the M-H subgroup, we identified significant enrichment of CSF1R:CSF1, CD70:CD27, TNFRS9:TNFS9, CTLA4:CD86/CD80 and CD28:CD86/CD80 receptor-ligand pairs (**Fig. 3B**). The higher expression of the CSF1R:CSF1 pair in the M-H subgroup is consistent with the M2 macrophage enrichment in this subtype and nominates a therapeutically actionable immune checkpoint axis for future translational studies guided by matrisome-based stratification of GBM patients. In conclusion, these observations indicate that the characteristics of the M-H TIME support a unique set of cellular and molecular features consistent with immune suppression.

**Figure 3.**
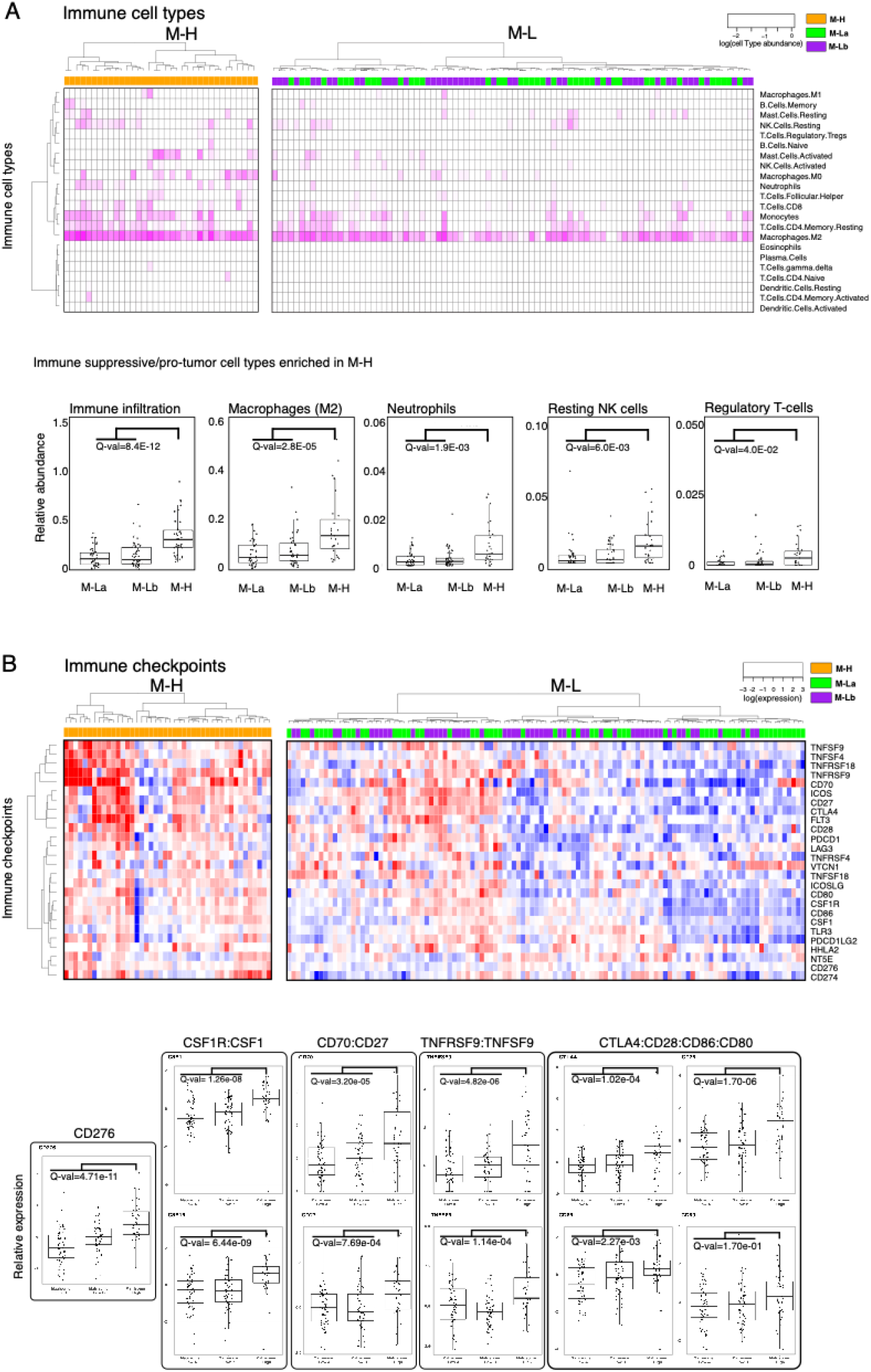
The GBM tumor-immune interactions across matrisome subtypes. ***A.*** *Enrichment of immune cell types in the microenvironment of M-H vs M-L tumors. Immune cell type fractions were quantified through CIBERSORT analysis. The immune infiltration level in each sample is quantified based on the methylation-based immune fraction scores* {Thorsson,2018, #287}*. The enrichment of each immune cell type is quantified through the multiplication of the total immune infiltration score and fraction of the corresponding immune cell type. The statistical significance is quantified through the Wilcoxon-signed ranked test. The resulting p-values for M-H vs. M-La and M-H vs. M-Lb are merged using the Stouffer method and corrected for multiple hypothesis testing across all immune cell types using the Bonferroni method. The immune cell type levels with significant differences across groups are shown in the box plots. **B.** Distribution of immune checkpoint expression across M-H vs. M-L tumors. We analyzed the mRNA expression of immune checkpoint molecules for which targeting strategies are in clinical use or trials* {Bozorgui, 2021 #505} *{Qin, 2019 #504}. The statistical procedure is identical to that in Figure 3A. The immune checkpoint axes within which both the receptor and the ligand are statistically different between the groups are shown in the box plots except for the orphan receptor CD276*.

### CMP-encoding genes are differentially expressed in vascular structures

To identify how matrisome varies across anatomical niches of GBM, we analyzed the localization and heterogeneity of the CMP-encoding gene expression. The analysis of mRNA expression in 245 samples from 7 anatomical regions across 34 tumors in the IvyGap dataset permitted spatial mapping of gene expression to distinct anatomic/histologic domains, including leading edge (LE, n=16) infiltrative (IFT, n=21), cellular tumor (CT, n=104), microvascular proliferation (MVP, n=22), hyperplastic blood vessels (HPBV, n=20), pseudopalisading cells around necrosis (PAN, n=39) and perinecrotic zone (PNZ, n=23) {Puchalski, 2018 #49}. Through a supervised hierarchical clustering analysis, we identified distinct gene expression clusters localized in vascular (MVP/HPBV) and infiltrative tumor-brain regions (IFT/LE) with a less distinct enrichment in regions associated with necrosis (PAN/PNZ) or solid cellular tumor (CT) without vascular or necrotic features (**Fig. 4A**). Although marked heterogeneity in the CMP-encoding gene expression is evident in samples from individual patients, the patterns of CMP-encoding gene expression within similar anatomic regions are largely conserved across patients (**Supplementary Fig. 4A**). This finding indicates that intratumoral matrisome heterogeneity is primarily driven by the differential expression across distinct anatomic regions.

**Figure 4.**
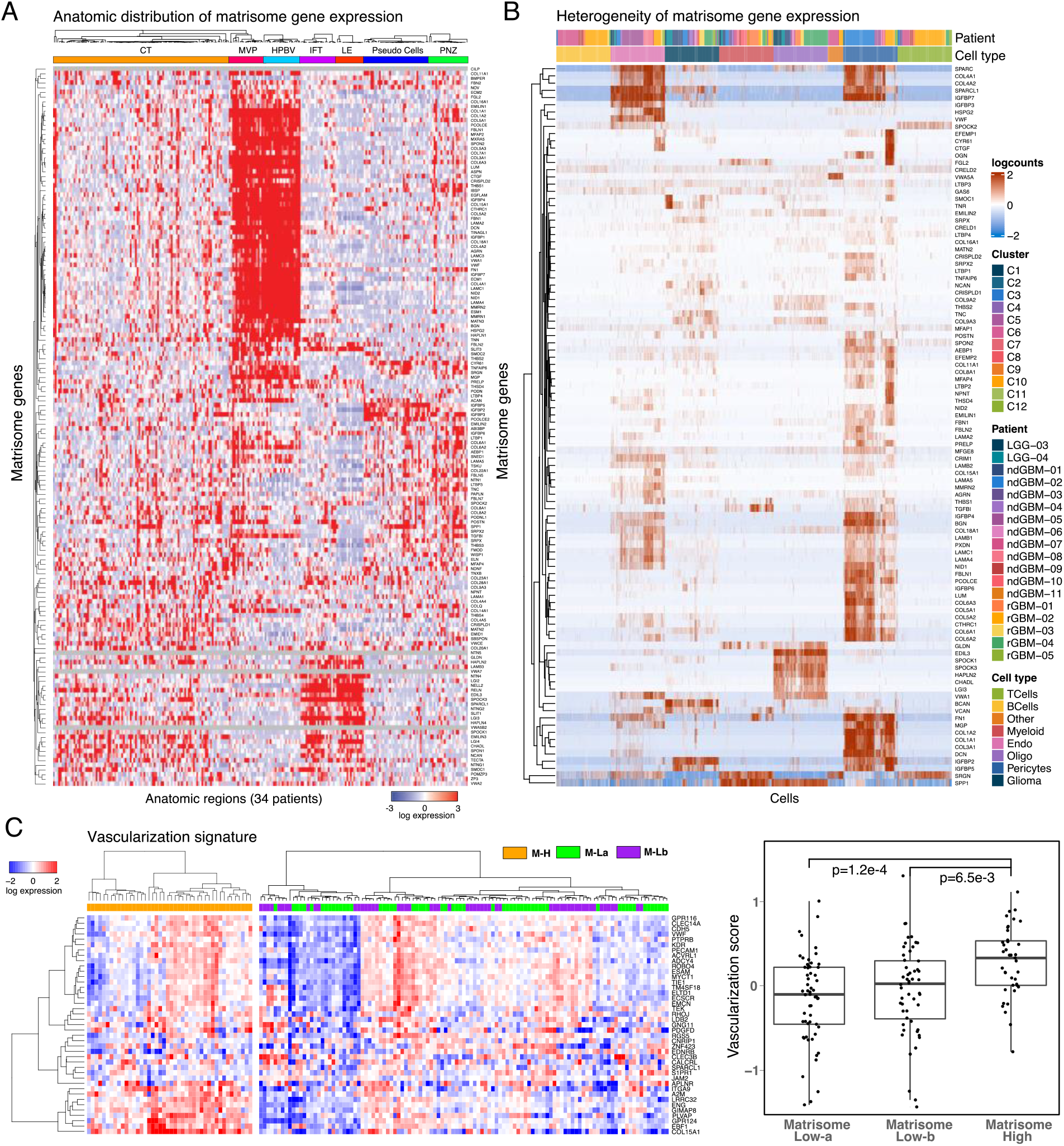
The spatial compartmentalization and heterogeneity of matrisome expression. ***A**. Supervised hierarchical clustering of anatomically resolved matrisome gene expression across 245 RNAseq samples in 34 patient tumors. **B.** The heterogeneity of matrisome gene expression based on single cell RNA transcriptomics (GSE182109) {Abdelfattah, 2022 #100}. **C.** The heatmap representation of the vascularization gene signature expression in M-H vs. M-L tumors (left) and the quantitative comparison of the vascularization in M-H vs M-La and M-Lb based on the vascularization score as defined in {Masiero, 2013 #348}. The p values are based on a non-parametric Wilcoxon test*.

We studied the specific GBM cell types underlying the CMP-encoding gene expression patterns using a large single cell RNA sequencing dataset with cell type annotations {Abdelfattah, 2022 #100} (**Supplementary Table 4**). Consistent with the localization in vascular regions, CMP-encoding gene expression is markedly increased in pericytes and endothelial cells, while expression level is modest in glioma cells and negligible in myeloid, B or T cell clusters (**Fig. 4B**).

A unique gene expression cluster is also identified in cells with oligodendrocyte markers (**Fig. 4B**); interestingly, genes in this cluster are highly expressed in LE/IFT samples, moderately expressed in CT, but virtually non-expressed in MVP, HPBV, PAN and PNZ samples according to the analysis of the IvyGap dataset (**Supplementary Fig. 4B**). To further examine the relationship between CMP expression and tumor vascularity, we analyzed vascularization markers in GBM samples (N=157) from the TCGA repository using a previously reported cancer vascularization signature {Masiero, 2013 #348}. The analysis identified enrichment of vascularization markers, and consequently suggested higher degree of vascularization, in the M-H vs M-La and M-Lb subgroups (**Fig. 4C**). In summary, the matrisome genes are predominantly expressed in the vascular structures and, in turn, associated with increased vascularization in GBM tumors.

### A matrisome signature establishes a prognostic marker for GBM

Given robust associations of the M-H subgroup with patient outcomes and tumor phenotypes, we sought to generate a minimal M-H specific gene set as a potential biomarker for GBM prognosis and treatment responses. Guided by a Lasso regression that identified most discriminating genes between the matrisome subgroups, we selected 17 genes highly enriched in the M-H subgroup (**Supplementary Fig. 5A**). With hierarchical clustering, we defined three distinct clusters across the TCGA GBM cohort (N=157) with high, intermediate, and low expression of these 17 genes (**Fig. 5A**) and designated them as M-H’ (N=32), M-I’ (N=55) and M-L’ (N=71), respectively. Importantly, these subgroups retained prognostic significance for OS and PFS, with M-H’ having the worse outcomes (log-rank p= 4.122E-3 and 5.392E-4, respectively). Moreover, the inverse relationship between the signature expression and survival is monotonous across the subtypes such that PFS and OS for M-H’, M-I’ and M-L’ were 3.9, 5.3, and 8.4 months and 10.4, 12.5, and 14.9 months, respectively (**Fig. 5B**). In summary, the 17-gene signature, which we referred as the “matrisome signature”, provided a refined prognostic marker for GBM.

**Figure 5.**
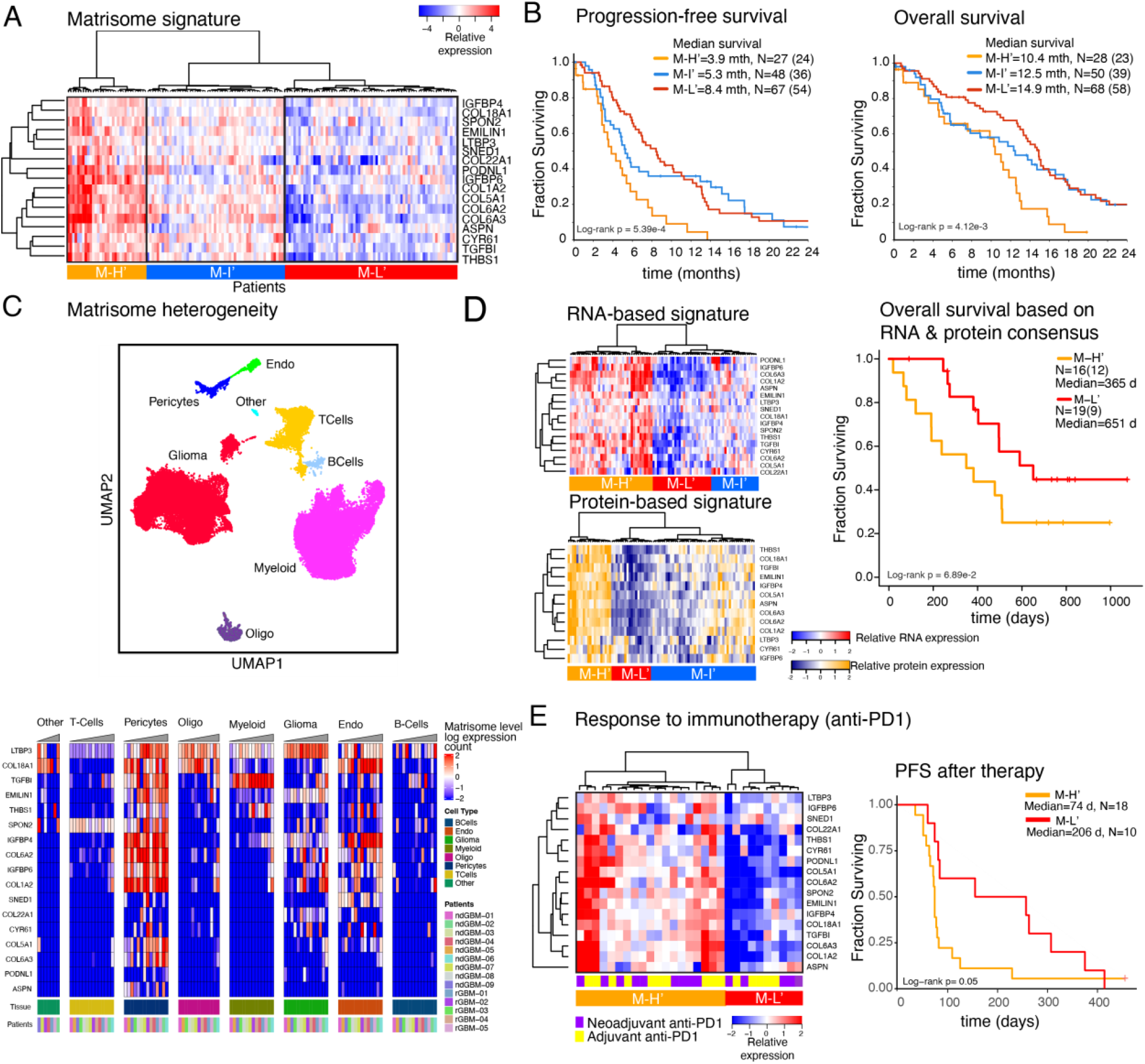
Predictive matrisome signature and response to immunotherapy. ***A.*** *The heatmap representation of the 17-gene matrisome signature based on the LASSO analysis to capture the most discriminant genes across the matrisome subtypes. 17-gene matrisome signature further refines the M-H’ cohort and resolves the M-La and M-Lb into matrisome Intermediate (M-I’) and low (M-L’) cohorts. **B.** The overall and progression free survival of patients with M-H’, M-I’, and M-L’ GBM as defined in (A).* (mth=months, truncated at 24 months). N is the number of patients. *The numbers in parenthesis designate number of events (i.e., progression for PFS and death for OS). **C.** The single cell transcriptomic analysis of the 17-gene matrisome signature. The distribution of cell types in the GBM ecosystem (top, adapted from* {Abdelfattah, 2022 #100}*) and the distribution of the matrisome signature expression across cell types (bottom) based on single cell transcriptomics. **D.** Transcriptomic (top, left) and proteomic (bottom, left) expression profiles of the matrisome signature in the matched tumor samples within the CPTAC repository. The proteomic analysis involves the 13 proteins that are encoded by genes within the 17-gene signature and are presented in the CPTAC mass spectroscopy data {Wang, 2021 #16}. The comparison of overall survival between the M-H’ vs. M-L’ patient cohorts (right). The M-H’ and M-L’ groups in the survival analyses represent the patients whose signature expression levels are concordant across the protein and RNA profiles.* d=days) ***E.*** *Matrisome-signature levels correlate with responses to PD1 blockade. The heatmap representation of the mRNA expression in tumor samples from recurrent GBM patients in the clinical trial with adjuvant and/or neoadjuvant anti-PD1 therapy {Cloughesy, 2019 #351} (left) and the PFS (from day of registration to progression per iRANO criteria or death) in M-H’ vs M-L’ patients that are stratified based on the matrisome signature expression* (d=days) (right).

Next, we characterized the heterogeneity and proteogenomic landscape of the matrisome signature. We mapped the heterogeneity of the matrisome signature through a single cell transcriptomics analysis (**Fig. 5C**). Consistent with the enrichment of the expression of CMP-encoding genes in the vascular niche, each gene in the signature is expressed at high levels in pericytes and/or endothelial cells, with a few exceptions (e.g., COL22A1). Although at less substantial levels, there is also variable co-expression of these 17 genes in other cell types, most predominantly in glioma cells. We investigated the proteogenomic reflection of the matrisome signature using transcriptomic and mass spectroscopy-based proteomic data from matched samples of GBM patients (CPTAC GBM database, N=86 after filtering out IDH-mutated and/or poor-quality samples based on pathology assessment) {Wang, 2021 #16}. Similar to the analysis of the TCGA-cohort, we observed three clusters in both proteomic and transcriptomic analysis that corresponded to M-H’, M-I’, and M-L’ subgroups. The matrisome signature is highly associated with poor survival, particularly for patients whose tumors are in the consensus M-H’ subgroup (i.e., highly expressed signature at both protein and RNA levels, see Methods) compared to the patients with M-L’ tumors (median survival times, M-H’: 365 days; M-L’: 651 days, log-rank p=0.069, **Fig. 5D**). Although the sample size is substantially smaller compared to the TCGA cohort, the analysis of the mRNA and the protein data individually also demonstrate strong trends of poor survival outcomes when the matrisome signature is expressed (**Supplementary Fig. 5D**). The results from the proteogenomic and single cell transcriptomic analyses are consistent with those from TCGA- and IvyGap-based gene-expression analyses for the patient subgrouping (M-H’, M-I’, M-L’), survival outcome trends, and anatomic distribution of the matrisome signature expression.

### Matrisome status correlates with response to immunotherapy in GBM

We asked whether the matrisome gene signature correlates with responses of GBM tumors to immunotherapy. We focused on a cohort of patients who carried recurrent and surgically resectable GBM, were profiled for mRNA expression, and received neoadjuvant and/or adjuvant anti-PD1 therapy (Ivy Foundation Early Phase Clinical Trials Consortium) {Cloughesy, 2019 #351}. The survival time covered the period from trial registration prior to therapy and surgery to second progression or death, respectively for patients with resectable recurrent disease. The unsupervised clustering of mRNA levels of the 17 matrisome genes partitioned the patients into M-H’ (N=19) and M-L’ (N=10) groups **(Fig. 5E)**. After anti-PD1 therapy, the patients with M-H’ tumors had significantly shorter PFS and OS than patients with M-L’ tumors (**Fig. 5E**, **Supplementary Fig. 5**, log-rank p-value = 0.05 for both PFS and OS). Patients who received the neoadjuvant plus adjuvant treatment and the adjuvant-only treatment were similarly distributed between groups, 9 out of 19 patients with M-H’ tumors and 5 out of 10 patients with M-L’ tumors received the neoadjuvant plus adjuvant treatment. Survival analyses therefore were unlikely to be confounded by the treatment difference between these two groups of patients. Next, we analyzed the survival outcomes in a control cohort (GLASS consortium) to rule out the possibility that the survival differences are merely a reflection of the prognostic impact of the matrisome signature independent of the anti-PD1 treatment {Varn, 2022}. The control cohort involved patients with recurrent and surgically resectable GBMs who were treated with chemotherapy and/or radiotherapy but not anti-PD1 therapy. In the control cohort, we focused on the patients with mRNA expression profiling after second surgery (i.e., recurrence) (N=164). First, we classified patients based on the matrisome signature status through unsupervised hierarchical clustering (M-H’, N= 86 vs. M-L’, N= 78) (**Supplementary Fig. 5**). Next, we analyzed the survival period from recurrence to death, a period equivalent to trial registration to death period for the anti-PD1 treated patients with recurrent disease. The survival difference between patients carrying M-H’ vs. M-L’ is more significant in the anti-PD1 treated cohort compared to that in immunotherapy-naïve control cohort (Hazard Ratio (HR) _anti-PD1_=0.38, HR_control_=0.85; log-rank P _anti-PD1_ = 0.05, log-rank P_control_ = 0.32). The comparison of the anti-PD1 trial cohort with the control cohort suggests that the matrisome status correlates with anti-PD1 treatment beyond the prognostic impact of the matrisome expression in patients with recurrent disease. The correlation between the patient survival, anti-PD1 treatment and matrisome status also justifies future clinical trials involving well-controlled study arms including anti-PD1 treated vs. untreated cohorts, and different matrisome states. Such future trials may establish an CMP-based clinical marker that guide selection of GBM patients who are likely to benefit from immunotherapy.

## Discussion

We report the clinical and phenotypical significance of CMP-encoding genes in IDH WT GBM. Expression profiles of matrisome genes and proteins in GBM have been reported {Laurentino, 2022 #349; Sethi, 2022 #350} but their biological and clinical relevance have not been well characterized. Using multi-modal molecular and clinical datasets from diverse sources (TCGA, IVYGap, CPTAC, GLASS, and GBM single-cell transcriptomics), we identify novel anatomic and cell-type specific CMP-encoding gene expression signatures that correlate with molecular subtypes and mutations, mesenchymal and immune TME phenotypes, survival, and potentially, responses to immunotherapy. These studies provide new insight into mechanistic roles of regionally expressed CMPs, which may serve as novel therapeutic targets and clinical biomarkers to stratify patients for prognosis and treatment response.

In our analyses, the M-H subgroup consistently emerges as the most robust indicator of poor survival outcomes, malignancy, and mesenchymal phenotypes. Although the M-H subgroup is enriched in the mesGBM transcriptional subtype, M-H predicts shorter survival even within the mesGBM subtype (**Fig. 1D**), indicating that it has clinical and functional significance independent of this association. The M-H subgroup is also associated with NF1 mutations {Verhaak, 2010 #293}, expressions of EMT-related genes and proteins {Zarkoob, 2013 #443}, increased infiltration of pro-tumor immune cells, including M2 macrophages and Treg cells {Chen, 2018 #370; Kaffes, 2019 #369}, and tumor angiogenesis/vascularization {Phillips, 2006 #218}. EGFR mutations and RTK pathway activation are enriched in the M-Lb subgroup, which has the best overall prognosis, while the M-La subgroup is enriched in the proneural GBM subtype **(Fig. 1-2 and Table 1).** Aside from the enrichment of the RTK pathway in M-Lb, very few significant associations between either of M-L subgroups and any specific proteomic signaling or immune cell phenotypes are detected. At the protein level, fibronectin, PAI1 and Caveolin 1 are upregulated in M-H and beta-catenin is upregulated in the merged M-L subgroup (M-La + M-Lb) **(Fig. 2B).** As noted above, Fibronectin, PAI1 and Caveolin1 are associated with increased GBM malignancy {Seker, 2019 #332; Kabir, 2022 #353; Wu, 2022 #352; Pu, 2019 #357; Moriconi, 2021 #355}. Contrary to the reported association of Wnt-signaling with mesenchymal GBM, we find that the Beta-catenin expression is reduced in M-H GBMs {Park, 2019 #360}. Together these observations indicate the unique significance of the M-H subgroup in promoting GBM malignancy related to mesenchymal, immune, and vascular phenotypes.

The robust correlations between M-H and immune features suggest functional roles for M-H in modulating an immune suppressive TME. As shown for mesenchymal GBM, M-H tumors have increased overall immune cell infiltration, particularly M2 macrophages, neutrophils, resting NK cells and Tregs {Chen, 2020 #368; Kaffes, 2019 #369; Chen, 2018 #370}. M2 macrophages and Tregs are implicated in immune evasion in part through immune checkpoint-mediated signaling and suppression of CD8^+^ cytotoxic T lymphocytes {Tu, 2021 #374}. The increase in resting versus activated NK cells may indicate a defect or reduction in innate immune responses {Wu, 2020 #375}. The increase in tumor-associated neutrophils (TANs) in M-H is consistent with the observed increase in neutrophils reported in MesGBM sub-type {Wang, 2017 #295}. Increased TANs generally predict poor outcomes, but their dual anti-tumor and immunosuppressive functions indicate that further study is required to determine the functional importance of increased TANs in M-H GBM {Shaul, 2019 #387}. Increased peripheral neutrophil:lymphocyte ratios predict shorter GBM patient survival {Chim, 2021 #379} and intratumoral neutrophils promote GBM malignancy in part through S100A4-mediated activation of glioblastoma stem cell (GSC) proliferation, invasion, and resistance to anti-VEGF therapy {Liang, 2014 #376}. Notably, S100A4 is a marker and a regulator of GSCs in a mouse model and is a master regulator of mesenchymal transitions in GBM cells {Chow, 2017 #378} and has recently been identified as a regulator of immune suppressive myeloid and lymphoid cells {Abdelfattah, 2022 #100}. Of interest, analysis of M-H DEGs reveals coordinated enrichment of phenotypes related to immunity and EMT, which indicate a potential role in orchestrating functional interactions between mesenchymal changes and the TME (**Fig. 2C**). This is consistent with the recently recognized correlations between mesenchymal GBM subtypes and distinct immune suppressive TMEs {Hara, 2021 #149; Wang, 2017 #295} and the robust reciprocal cross talk between EMT and tumor immune landscapes identified in other cancers {Reiman, 2010 # 366; Taki, 2021 #363}.

Overall, expression of specific immune checkpoint genes and many receptor-ligand pairs implicated in GBM malignancy and immune suppression are increased in M-H GBMs, including CD276 {Zhang, 2018 #380}, CSF1R:CSF1 {Pyonteck, 2013 #381}, CD70:CD27 {Wischhusen, 2002 #382}{Jin, 2018 #383},TNFRSF9:TNFSF9 {Freeman, 2020 #384}{Cho, 2021 #385}, CTLA4:CD80/86, and CD28:CD80/CD86 {Guan, 2021 #386} **(Fig. 3B).** Challenges exist in deconvoluting the state of immune suppression, which depends on complex and often antagonistic effects that are dependent on not only cell composition and immune checkpoint expression but also specific cell-cell interactions and systemic factors. Nevertheless, the data collectively supports a conclusion that the M-H phenotype is more immune suppressive than its M-L counterparts. A potential consequence of a more immune suppressive TME is the resistance to immune checkpoint therapy. The conclusion that M-H subgroups are more immune suppressive is bolstered by our finding that responses to anti-PD-1 blockade are limited in GBMs with high expression of the M-H gene signature **(Fig. 5E).**

The spatial analysis using the IVYGap dataset shows that the strongest matrisome enrichment is in vascular structures (MVP and HPBV) and the LE/IFT regions at the tumor margins **(Fig. 4A).** These domains have been identified as distinct niches that harbor and maintain GSCs {Fidoamore, 2016 #445; Hide, 2018 #446; Jung, 2021 #444}, suggesting that CMP “codes” may define specific anatomic and functional domains in GBM. Given the central role of GSCs in GBM treatment resistance, progression, and dissemination, CMPs may promote oncogenic signaling and treatment resistance in GSC supportive niches. This new understanding of niche-specific CMP expression profiles is expected to enable rational targeting of matrisome protein-GSC mechanisms in future studies. Consistent with CMP enrichment in the vascular niche, single cell transcriptomic analysis identified pericytes and endothelial cells as primary contributors of CMP expression with modest contributions from glioma cells and oligodendrocyte type cells **(Fig. 4B).** The enrichment of a prognostic ECM-signature expression in pericyte niches within recurrent GBM tumors has also been reported in a recent study {Hoogstrate, 2023}. Among immune cells, CMP-encoding gene expression is negligible in B and T cells and is slightly higher but still sparsely expressed in myeloid cells **(Fig 4B).** The oligodendrocyte-enriched cluster localizes primarily within the LE/IFT regions **(Supplementary Fig. 4),** suggesting potential unique roles in promoting GBM invasion.

The associations between mesenchymal phenotypes, angiogenesis, tumor microenvironment and immune infiltrates have previously been reported {Phillips, 2006 #218} {Chen, 2018 #370; Tian, 2020 #455; Varn, 2022}. The strong associations of such key oncogenic processes with CMP enriched signatures which we report here, suggest that CMPs function as a structural and functional hub critical to the integration of oncogenic crosstalk in the vascular niche. The relevance of matrisome in regulating angiogenesis and immune stroma of the TME, and modulating responses to genotoxic and targeted drugs as well as immunotherapies have been studied in the context of diverse solid tumor types {Henke, 2019 #56}, {Pietila, 2021 #328} {Bin Lim, 2019 #327; Yang, 2021 #14}. The results from these studies further support our findings in GBM.

This study identifies a novel matrix code in GBM that correlates with patient survival as well as response to immunotherapy and correlates with hallmark features of mesenchymal change, immune suppression, and tumor angiogenesis. The matrix code is enriched in specific GBM niches, particularly in functional vascular and infiltrative domains central to GSC localization and maintenance. More detailed and precise mapping of CMP-cellular networks in specific GBM niches will inform critical signaling hubs whose functional and therapeutic importance can be further validated in *ex vivo* experimental 3D models. In addition, the relevance of a 17-gene matrisome signature for survival and immunotherapy responses indicates a potential for the development of new clinical biomarkers. The prognostic and predictive value of this matrisome signature warrants validation in future clinical trials where it may prove to be a useful guide for individualized patient management. We did not identify any specific CMP signatures in hypoxic niches, which have great significance for treatment resistance, GSCs and immune suppression. Future studies may be required to deconvolute these expected interactions with the matrisome. Future immunohistochemical and molecular spatial studies can define the broader role of the matrisome in GBM through analysis of matrisome associated proteins. A key future direction is to develop therapeutic strategies tailored to matrisome composition. Such therapeutic strategies may disrupt matrisome-mediated oncogenic signaling and thus sensitize tumors to immunotherapy or other therapeutic modalities to improve survival outcomes of patients with this devastating disease.

## Author Contributions

AK, MV, ZD and RCR conceptualize the study, RCR, MV and AK prepare the manuscript, AK, ZD, XL, BB (TCGA database analysis), MV (IVYGap GBM), ZY (multivariate analysis), MK, AK, KT, and OB (proteomics), NA, EKK, AK (scRNAseq), KY supervise the scRNAseq analysis, RCR and AK supervise the study.

## Acknowledgements

This study was supported by grants John S. “Steve” Dunn, Jr. & Dagmar Dunn Pickens Gipe Chair in Brain Tumor Research Foundation (# 20090011, 2019 SPG Performance grant and NIH funded R01-CA181445. We are also thankful to the Departments of Bioinformatics and Computational Biology, at MD Anderson Cancer Center, Houston, TX USA. This work is supported with grants from MDACC Support Grant P30 CA016672 (the Bioinformatics Shared Resource) (AK), U01CA217842 (AK). This manuscript was edited at Life Science Editors.

## Data availability statement

All the data sets used in this manuscript are publicly available. The accession codes and links to repositories are provided in methods or other relevant sections.

## Code availability statement

The pieces of code and scripts which were used to execute the analyses are available upon request.

## Methods

### TCGA datasets-based multi-omic analyses

The TCGA datasets-based genomic, transcriptomic, and proteomic analyses were performed using data available from cBioportal (https://cbioportal-datahub.s3.amazonaws.com/gbm_tcga_pub2013.tar.gz, https://cbioportal-datahub.s3.amazonaws.com/gbm_tcga.tar.gz) {Hoadley, 2018 #484; Brennan, 2009 #116; Cerami 2012}}. For each data modality IDH1 mutations were filtered out. The transcriptomic analysis included RNA expression data from 157 tumor samples of 151 GBM patients. The RNA expression analysis of matrisome protein-encoding genes (**Fig. 1B**) was initiated with 274 genes. The genes with low standard deviation across samples (σ<1) were filtered out to enable classification of patients with more variant and likely discriminant transcription events. In all analyses, hierarchical clustering of the RNA expression levels was performed using Manhattan distance and Ward method (heatmap.2 function in the gplot R package). Survival analysis was performed with the Kaplan-Meier method for censored data (Survival package in R in Figure 5) and cBioPortal survival analysis module (**Fig. 1**, **5B**) for group comparisons. The statistical significance of survival differences is evaluated with log-rank test. The enrichment analysis of matrisome subtypes across demographic groups and transcriptional subtypes were performed using a Fisher’s exact test. In each test, the enrichment of the most common matrisome subtype in a corresponding patient group (e.g., male) was tested against all other matrisome subtypes and other patient groups (e.g., female) using a 2×2 contingency table. The statistical difference of Karnofsky performance score and median age across matrisome subtypes were analyzed using a 1-way ANOVA test. The mutation data included whole exome sequencing data from 147 patients for which RNA expression data was also available. The mutational and copy number alteration oncoprints were generated using the oncoprint module in cBioPortal. The mutational and copy number enrichment analyses across matrisome groups were performed based on a Fisher’s exact test followed by Benjamini-Hochberg FDR-correction. The proteomics-based pathway scores were based on reverse phase proteomics data and the pathway score definitions and gene/protein lists in {Akbani, 2014 #339} (supplementary table 13 in the referenced article). In the referenced paper by {Akbani, 2014 #339}, members of each pathway had been predefined based on a Pubmed literature search on review articles describing the various pathways in detail. The batch corrected RPPA data had been median-centered and normalized across all samples to obtain the relative protein level. The pathway score was then the sum of the relative protein level of all positive regulatory components minus that of negative regulatory components in a particular pathway. We extracted the pathway scores for the IDH-wt GBM samples. The included pathways were apoptosis, cell cycle, DNA damage response pathway, core reactive pathway (stroma signature), EMT pathway, hormone receptor pathway, RAS/MAPK, EMT, mTOR/TSC. The transcriptomics based EMT scores were computed using the epithelial and mesenchymal gene signatures as defined in {Mak, 2016 #333}. The vascularization score was calculated using the gene signature defined in {Masiero, 2013 #348}.

### Multivariate analysis

For multi-variate analysis, a 156-by-6 data table was created by combining the features of CMP subgroups with two demographic features (Age and Gender), two disease phenotype features (GBM subtype and MGMT Methylation status) and one categorical feature for treatment type using Matlab R2020a Update 5, maci64. GLM was created through the fitglm function of Matlab using the 156-by-6 data table as input, and overall survival (in months) as response variable. The optional parameter of ‘distribution’ for the fitglm function was set as ‘poisson’ to be consistent with the characteristics of response variable, while default values were used for all other parameters. The 156-by-6 data table has five categorical variables and one continuous variable (Age), and fitglm picked a reference category for each of the five variables, i.e., M-H for CMP sub-group, Female for Gender, Classical for GBM Sub-type, Methylated for MGMT Methylation status and Chemoradiation for Treatment. The outputs of the fitglm included Coefficient Estimate (ranged [-1,1]), Standard Error, t-stat and p-value, and together they quantified the impacts on overall survival when each of the five categorical variables changed from the reference category to other possible values, with positive coefficient and t-stat indicating increased overall survival than the reference category. Meanwhile the same panel of outputs quantified the impact of increasing age to overall survival.

### Selection of genes contributing to matrisome subtypes

To select genes discriminating the matrisome subtypes, we performed multinomial logistic regression with Lasso regularization using the TCGA GBM dataset of mRNA expression. The mRNA expression was quantified by RSEM and followed by log2p1 transformation. In this analysis, 157 samples (44, 58, and 55 samples from M-H, M-La, and M-Lb subtypes, respectively) with 274 GBM core matrisome genes were included. The function cv.glmnet of the R package glmnet was used to do a 10-fold cross validation for selecting an optimal value from a series of regularization parameter λ’s. At the optimal λ, where the multinomial deviance is minimal, 47 genes were identified with non-zero coefficients from the multinomial regression model. Finally, from these 47 genes, we obtained 17 genes that were selectively and highly expressed in the M-H subtype and belonged to the same cluster in an unsupervised hierarchical analysis (**Supplementary Fig. 5B**).

### Immune cell infiltration and immune checkpoint analysis

The total leukocyte fraction and proportion of immune cell types for the GBM samples in the TCGA cohort were imported from {Thorsson, 2018, #287}. The methylation based total leukocyte fraction had been calculated as explained in {Hoadley, 2018 #484} and captured the leukocyte fraction based on the methylation probes with the greatest differences between pure leukocyte cells and normal tissue followed by estimation of leukocyte content with a mixture model. The proportion of immune cell types with respect to each had been inferred by a CIBERSORT analysis of transcriptomics data {Thorsson 2018, #287}. The absolute fractions of immune cell types in each tumor were estimated through multiplication of the proportion of the immune cell types in the immune cell population with total leukocyte fraction to enable cross sample comparisons of specific immune cell types. In total, immune cell infiltration from 153 samples (42 M-H, 58 M-La, 53 M-Lb) were analyzed. The levels of total infiltration and immune cell types were compared between the M-H and M-La/b groups using the Wilcoxon test. For an integrated M-H vs. M-L comparison, the p-values for “M-H vs. M-La” and “M-H vs. M-Lb” were merged with the Stouffer method adjusted for multiple hypothesis testing using the Bonferroni method. For immune checkpoint analysis, we curated 31 therapeutically actionable immune checkpoints, which have been subject of immunotherapy clinical trials {Bozorgui, 2021 #505}. The RNA expression levels of immune checkpoints were analyzed for the same patient cohort as before: 153 samples (42 M-H, 58 M-La, 53 M-lb). Excluded are those 4 samples from 2 patients with conflicting matrisome identities. The significance of differences was assessed using a Wilcoxon test followed by a Stauffer meta-analysis to merge the P-values for M-H vs. M-La and M-H vs M-Lb. The resulting p-values were adjusted for multiple hypothesis testing using Bonferroni method.

### IVYGap Analyses

Anatomic and histologic domains in which CMP gene expression was enriched were identified through analysis of the IVYGap database {Puchalski, 2018 #260}. IVYGap provides transcriptional signatures from laser capture dissected regions within GBM segmented by histologically defined anatomic structures or enrichment in putative cancer stem cell gene expression identified by RNA in situ hybridization. In this study we included only IDH wild type patients (n=34) and excluded IDH mutant patients. Therefore, the anatomic RNAseq study included 110 samples across 9 patient tumors and cancer stem cell RNA-seq study included 135 samples across 31 patient tumors. The anatomic domains include-Leading edge (LE) at the margin of tumor, infiltrating tumor (IFT) between leading edge and tumor core; and Cellular tumor (CT) regions comprising the tumor core. Within CT, subregions are identified based on structural features as follows: Hyperplastic Blood Vessels (HBV), Microvascular Proliferation (MVP), Pseudopalisading cells around necrosis (PAN) and Perinecrotic zone (PNZ). Since both reference histology and cancer stem cell sample sets were annotated by the above anatomic domains, they could be combined for regional analysis of correlations with CMP gene expression. Using the combined RNAseq data we performed unsupervised and supervised hierarchical clustering to identify enrichment of CMP gene expression and signatures in specific anatomic structures. Gene expression data was aligned against hg19 reference assembly, normalized (TbT normalization method as described {Kadota, 2012 #408}) and analyzed to quantitate FPKM value of each gene. Anatomic and cancer stem cell RNA-seq studies were selected to define structural features and performing hierarchical clustering using MORPHEUS software (https://software.broadinstitute.org/morpheus). RNAseq data from multiple samples within individual patient tumors was used to evaluate intra-tumoral heterogeneity of CMP gene expression.

### Single Cell RNAseq

Single-cell RNAseq data were mined from GSE182109 (https://www.ncbi.nlm.nih.gov/geo/query/acc.cgi?acc=GSE182109) {Abdelfattah, 2022 #55} and processed according to the authors’ previously described methods {Abdelfattah, 2022 #55}. Data were analyzed using Seurat V4.0.0 (RRID:SCR_016341) and ggplot2 V3.3.3 (RRID:SCR_014601) and ComplexHeatmap V2.7.8.100 (RRID:SCR_017270) packages were used for visualization.

### Proteogenomic analysis

Mass spectroscopy-based proteomics and RNA sequencing data available from the CPTAC glioblastoma repository {Wang, 2021 #16} (https://pdc.cancer.gov/pdc/study/PDC000204) were analyzed to classify patients based on the 17-gene matrisome signature gene expression. The tissue from 99 patients had been profiled with mass spectrometry analysis using the 11-plexed isobaric tandem mass tags (TMT-11). The RNA expression had been profiled from the matched samples by sequencing on HiSeq 4000 as paired end 75 base pairs. Excluded samples from our analysis were the cases with IDH1 hotspot mutations (6 patients), low-quality cases that failed the pathology evaluation (6 samples with low tumor nuclei, low cellularity, and high necrosis) and one case from a patient who died immediately after surgery due to intracerebral hematoma. The resulting 86 cases carrying IDH1 wild type tumors with high quality based on pathology review are used in the matrisome analysis. Proteomics data covered 13 of the 17 proteins (COL22A1, PODNL1, SNED1, SPON2 proteins were captured in less than half to none of the samples). The Z-scores of log-transformed protein expression levels are used for the analysis. The RNA expression data covered all the 17 genes in the signature. RNA expression data is median normalized across the samples and log-transformed. Hierarchical unsupervised clustering (Manhattan distance and Ward’s method) was applied to both RNA and protein expression data to select the patients with differential matrisome signature expression. The protein/RNA consensus M-H’ cohort which involved the patients with high matrisome signature expression at both RNA and protein level is identified. To define the consensus, first, the patients in the M-H’ cluster in the proteomics-based unsupervised analysis is included. Next, to eliminate the cases that are not concordant with RNA expression levels, we quantified a transcriptomic matrisome signature score for each sample as the sum of RNA expression levels across the signature genes (log normalized values as used in unsupervised clustering). For the M-H’ consensus we filtered out the samples with transcriptional matrisome scores levels below the upper 35% percentile. This filtering provides a group of samples that co-cluster in the high protein expression group and carry high RNA expression that could support the observed proteomic levels, meanwhile it eliminates the likely false positives in which the high protein expression is not backed by high RNA levels. We also included the select cases in the M-I’ group that carry both high mRNA (> 90 percentile) and high proteomic (the median signature protein expression level falls into M-H’ cohort range). Similarly, the consensus M-L cohort is identified to include matrisome low samples in protein levels only when matrisome RNA levels are also below average (lower 50%, Z<0) to exclude potential false negatives (i.e., low matrisome cases). We also included the cases in the M-I’ group that carry both low mRNA (< 10 percentile) and high proteomic (the median signature protein expression level falls into M-L’ cohort range). The overall survival for the M-H and M-L levels were compared using Kaplan-Meyer curves and log rank statistics with censored overall survival data.

### Analysis of response to PD1 blockade

The association of CMP gene expression with immunotherapy was investigated using pre-treatment RNA expression and PD1 blockade (with pembrolizumab) data from GBM patients. The mRNA analysis involved 29 patients treated with neoadjuvant or adjuvant anti-PD1 agent pembrolizumab. The RNA expression count data was imported from Gene Expression Omnibus under accession number GSE121810 (ncbi.nlm.nih.gov/geo/query/acc.cgi?acc=GSE121810). The clinical data available for 28 patients is based on the reference {Cloughesy, 2019 #351} and clinical trial (NCT02852655), and was kindly provided by Dr. Robert M Prins. The raw count data was subject to count per million count normalization and log transformation. Next, the log count normalized expression levels of the genes in the 17-gene matrisome signature were median normalized across the samples and analyzed with a hierarchical clustering (Ward method, Manhattan distance). The progression free and overall survival of patients in the matrisome high vs. low clusters were compared using the Kaplan-Meier curves based on the censored survival data (N=28). The significance of survival differences is assessed with a log-rank test. The median survival times based on Kaplan-Meier curves were reported (survival package in R). For the control group analysis, the transcriptomics and clinical (survival, recurrence, and surgery interval annotations) data for samples that were not treated with anti-PD1 therapy were downloaded from the GLASS consortium repository at https://www.synapse.org/#!Synapse:syn26465623 {Varn, 2022}. We included only the samples with transcriptome, overall survival, and time to second surgery (recurrence) were used (N=165). The TPM-normalized transcription data, which were from samples of patients after second surgery (recurrence) is analyzed with unsupervised hierarchical clustering (Ward method, Manhattan distance) to identify the M-H’ vs. M-L’ patients based on the 17-gene matrisome signature. The survival from recurrence to death is approximated through subtraction of the interval between the first (immediately after first diagnosis) and second surgery (at recurrence) from the overall survival. The recurrence to death survival times, log-rank P values, and Hazard rations between M-H’ vs. M-L’ patients were analyzed with Kaplan Meier curves using the Survival package in R.

## Supplementary figures

### A prognostic matrix code defines functional glioblastoma phenotypes and niches

**Supplementary Figure 1.**
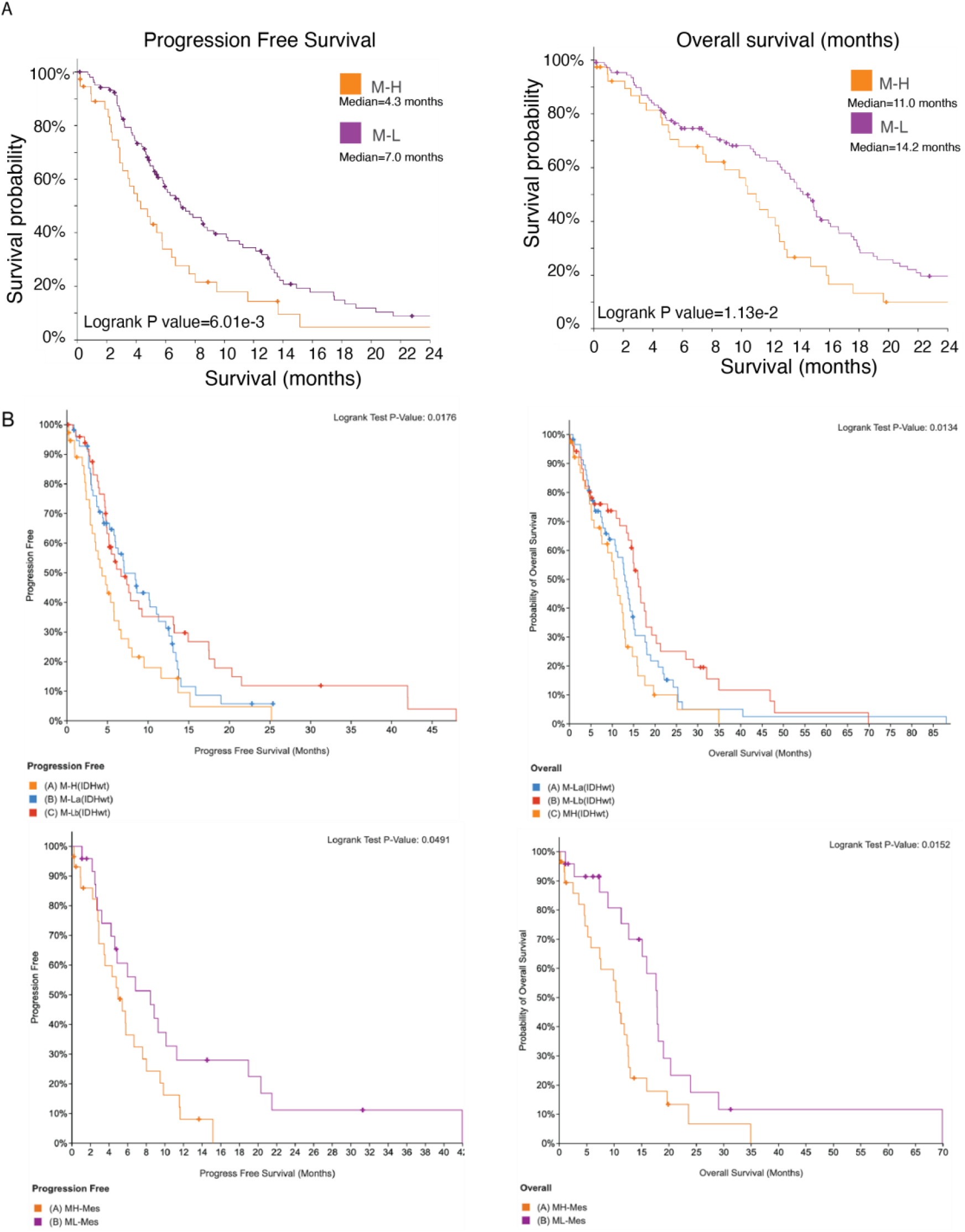
The landscape of ECM composition and patient survival in glioblastoma. ***A.** Kaplan-Meier analysis comparing overall and progression-free survival of the M-H subgroup with the M-La and M-Lb subgroups merged as M-L. **B.** The complete and untruncated survival curves for analyses in figure 1C and D. All underlying data and statistics are identical.*

**Supplementary Figure 2.**
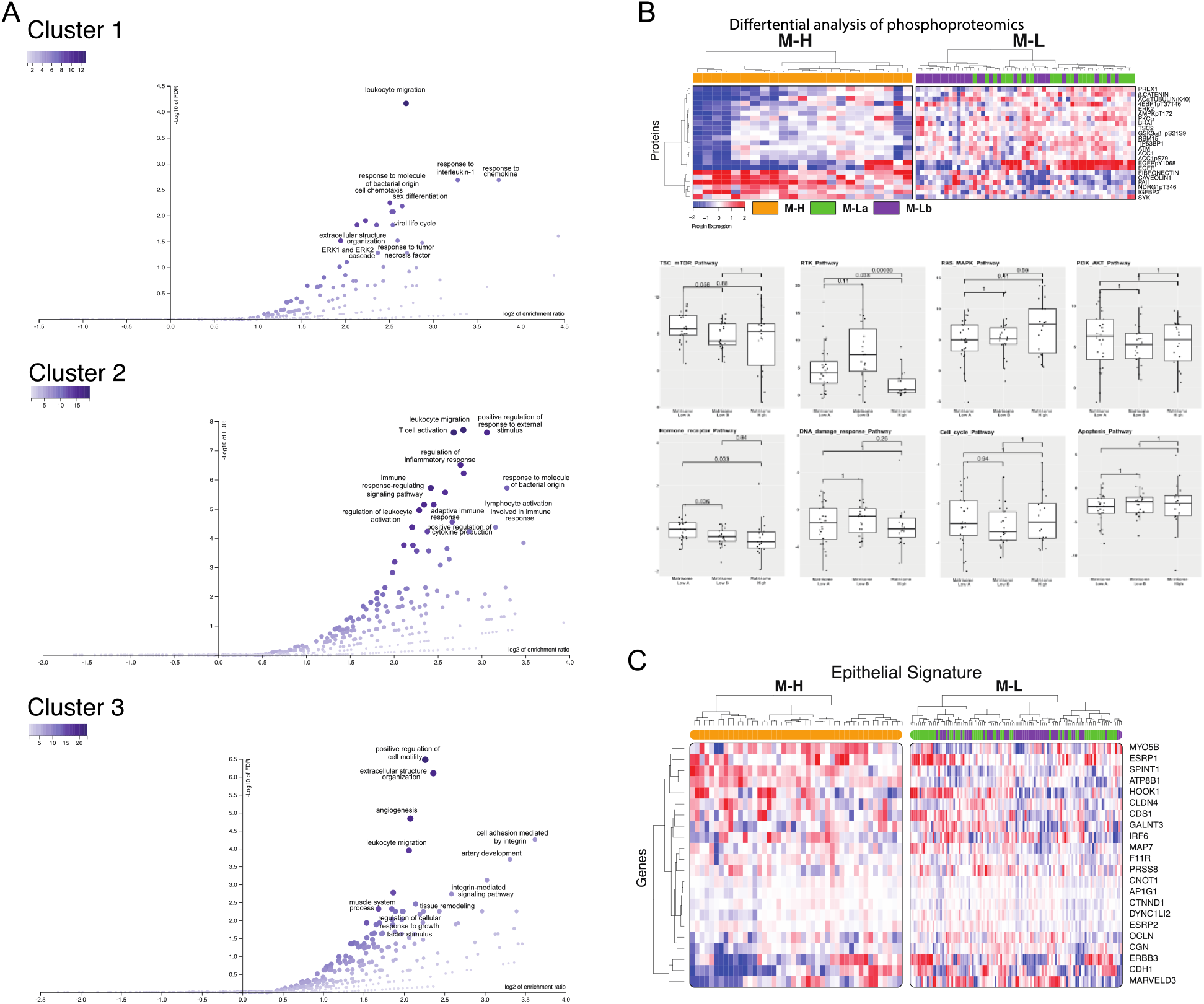
The differential multi-omic landscape of matrisome-enrichment in GBM. ***A.** Gene set enrichment analysis of each of the clusters in Figure 2B. **B.** The heatmap of phosphoproteomic entities with significant enrichment in M-H vs. M-L groups (top). The phosphproteomics-based pathway score differences between M-H, M-La and M-Lb groups. **C.** The transcriptomic analysis of epithelial signature (see main text figure 2 for the mesenchymal signature)*.

**Supplementary Figure 3.**
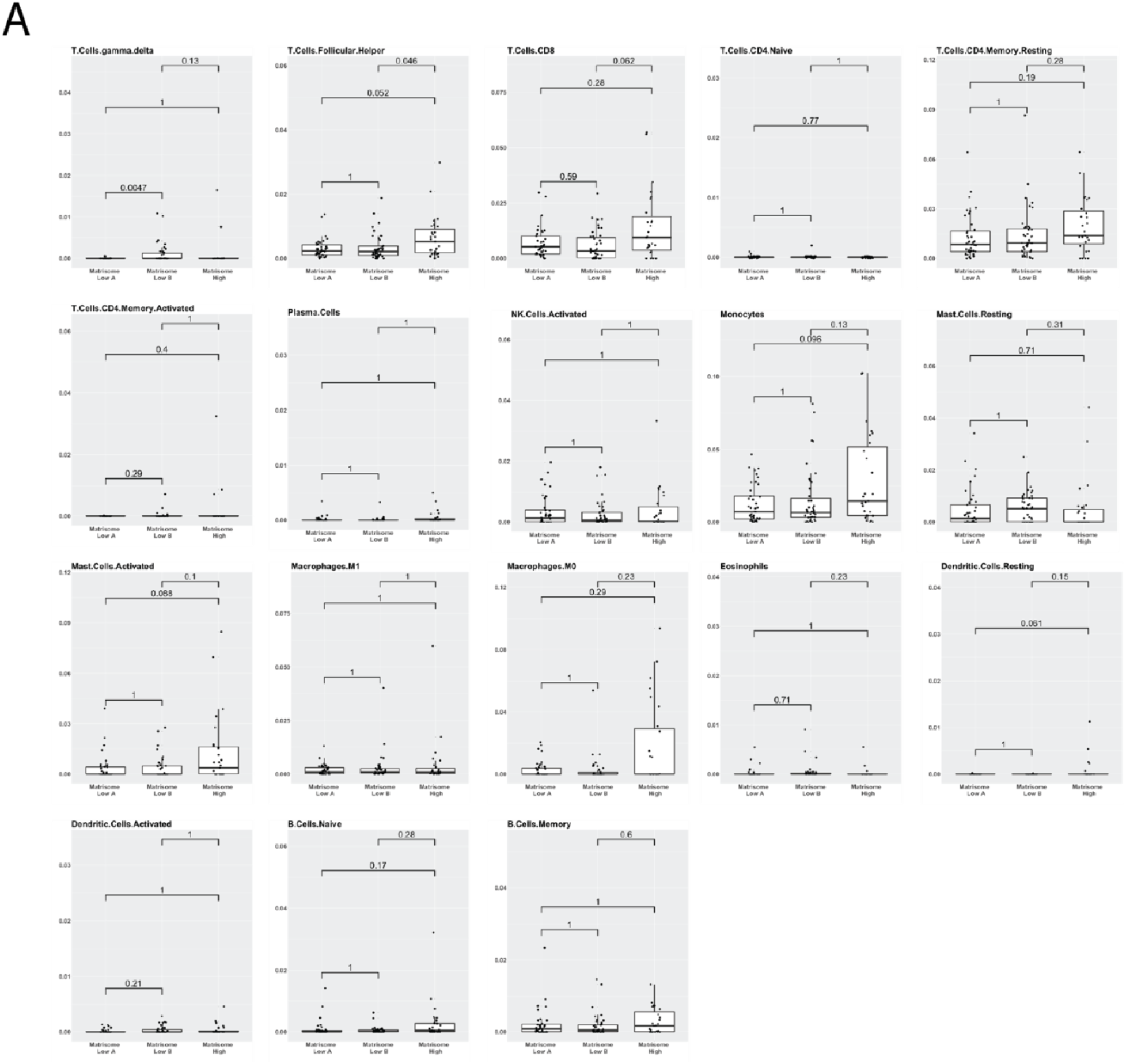
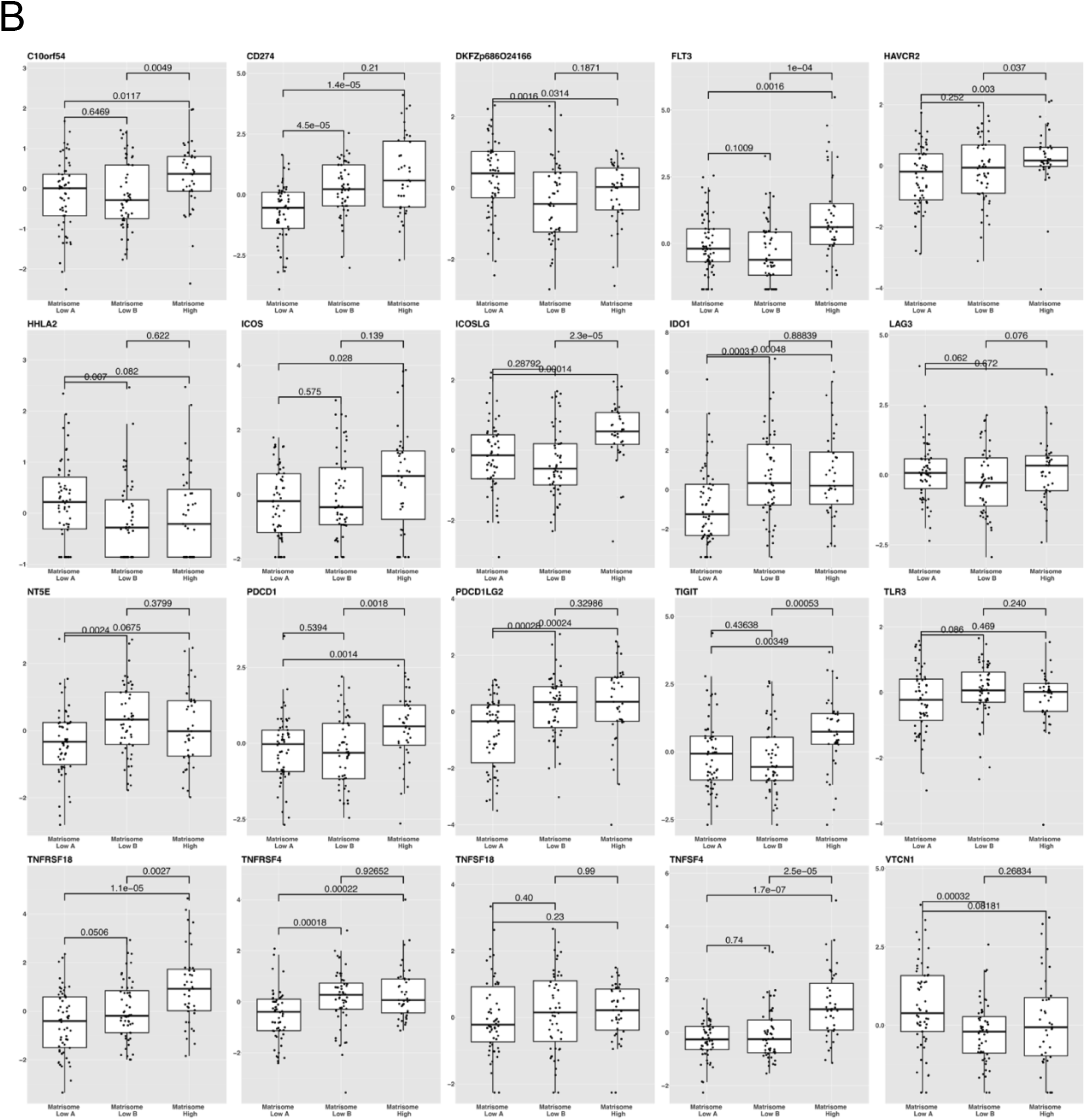
The GBM tumor-immune interactions across matrisome subtypes. ***A.** CIBERSORT-based analysis of immune cell type enrichment in M-H vs M-La and M-Lb groups. For statistically significant enrichments of cell types and overall immune infiltration, see figure 3A. **B.** The analysis of immune checkpoint expression in M-H vs M-La and M-Lb groups. For statistically significant enrichment of receptor-ligand pairs, see figure 3B.*

**Supplementary figure 4.**
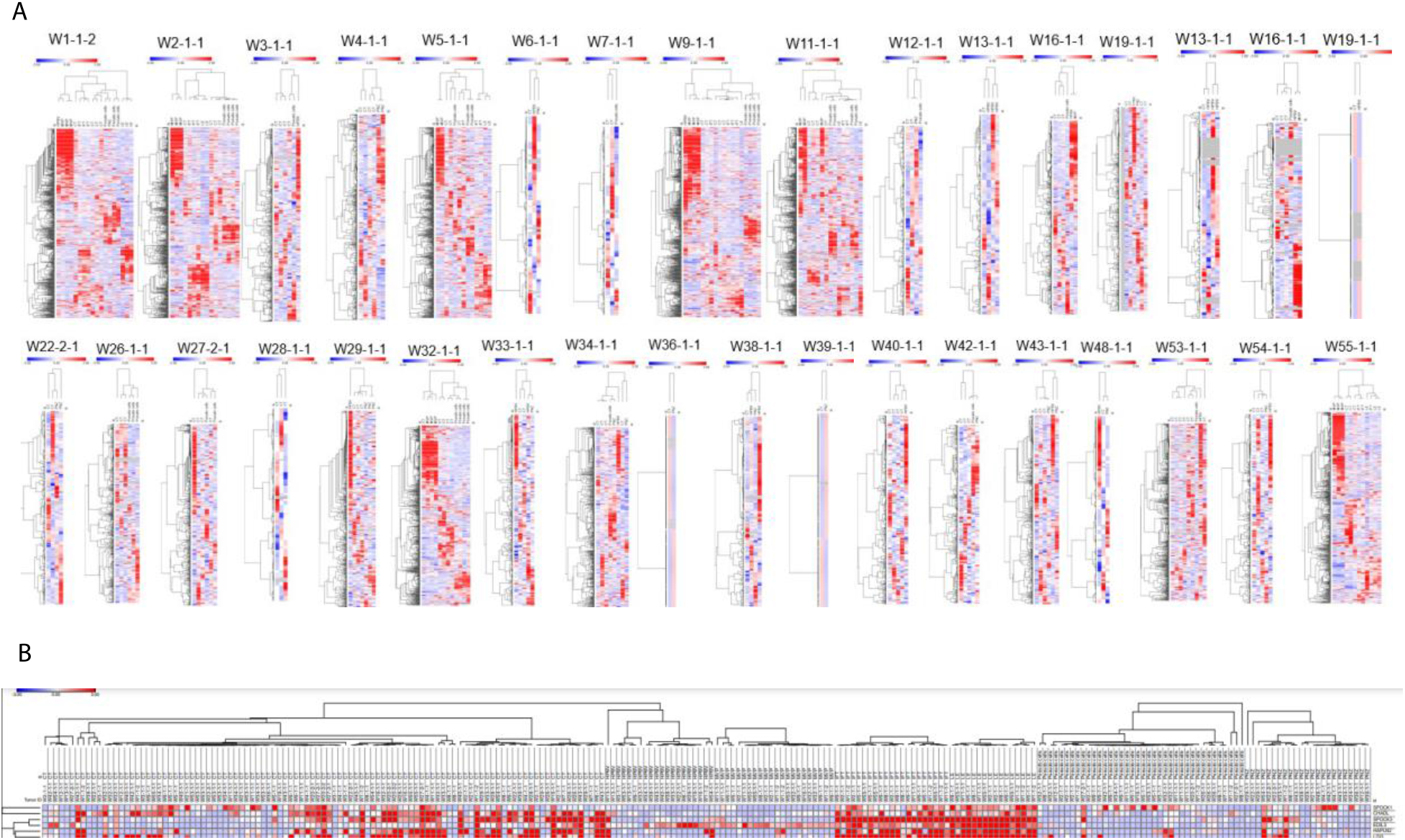
The spatial compartmentalization and heterogeneity of matrisome expression. ***A.** Hierarchical clustering analyses performed on multiple regional samples (Total=245 samples) of individual patients (n=34) show intratumoral heterogeneity based on anatomical regions. Each heatmap represents an individual patient. **B.** Supervised hierarchical clustering analysis of Oligo cell-type specific CMP genes.*

**Supplementary figure 5.**
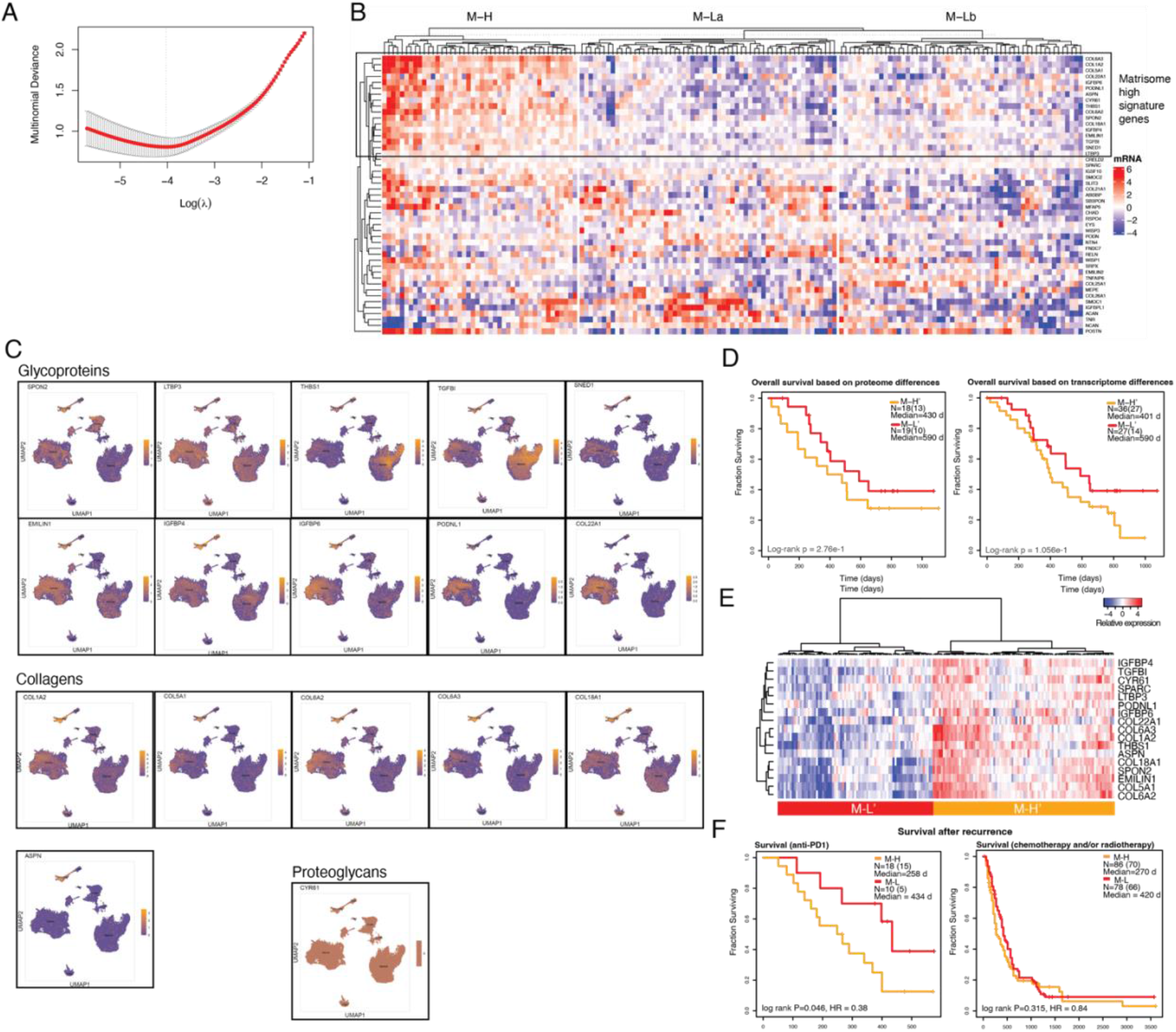
Figure 5. Predictive matrisome signature and response to immunotherapy. ***A.** LASSO analysis to identify most discriminate matrisome genes. Multimonial deviance as a function of log(λ) from cross validation. The red dots and error bars represent means and standard deviations. The vertical dotted line indicates the optimal log(λ) where the multinomial deviance is minimal. **B.** Heatmap showing the selected 47 genes with 157 samples from M-H, M-La, and M-Lb subtypes. The black frame emphasizes the 17-gene matrisome signature genes that are elevated in the M-H subtype. **C.** The single cell transcriptomic distribution of the genes across cell types in the matrisome signature as demonstrated through UMAP analysis. **D.** The overall survival of patients (Source: CPTAC, Wang et al, 2021) stratified as M-H’ vs. M-L’ based on the 17-gene matrisome protein (left) and mRNA (right) expression signature. **E.** The matrisome gene expression analysis of patients with recurrent/resectable GBM tumors not treated with anti-PD1 therapy (source: GLASS consortium repository, N=165). **F.** The recurrence to death survival period for patients treated with anti-PD1 therapy (left) and chemo/radiation therapy (GLASS consortium) (right).*

